# Evaluation of sequence-based tools to gather more insight into the positioning of rhizogenic agrobacteria within the *Agrobacterium tumefaciens* species complex

**DOI:** 10.1101/2024.04.18.590036

**Authors:** Pablo R. Vargas, Nuri Kim, Marc Venbrux, Sergio Álvarez-Pérez, Hans Rediers

## Abstract

Rhizogenic *Agrobacterium*, the causative agent of hairy root disease (HRD), is known for its high phenotypic and genetic diversity. The taxonomy of rhizogenic agrobacteria has undergone several changes in the past and is still somewhat controversial. While the classification of *Agrobacterium* strains was initially mainly based on phenotypic properties and the symptoms they induced on plants, more and more genetic information has been used along the years to infer *Agrobacterium* taxonomy. This has led to the definition of the so-called *Agrobacterium tumefaciens* species complex (Atsc), which comprises several genomospecies. Interestingly, the rhizogenic *Agrobacterium* strains are found in several of these genomospecies. Nevertheless, even up until today *Agrobacterium* strains, and in particular rhizogenic agrobacteria, are prone to misclassification and considerable confusion in literature.

In this study, we evaluated different phylogenetic analysis approaches for their use to improve *Agrobacterium* taxonomy and tried to gain more insight in the classification of strains into this complex genus, with a particular focus on rhizogenic agrobacteria.

The genome sequence analysis of 579 assemblies, comprising *Agrobacterium*, *Allorhizobium* and *Rhizobium* strains demonstrated that phylogenies based on single marker genes, such as the commonly used 16S rRNA and *recA* gene, do not provide sufficient resolution for proper delineation of the different genomospecies within the Atsc.

Our results revealed that (in silico) multi-locus sequences analysis (MLSA) in combination with average nucleotide identity (ANIb) at a 94.0% threshold delineates genomospecies accurately and efficiently. Additionally, this latter approach permitted the identification of two new candidate genomospecies.

## Introduction

The genus *Agrobacterium* is part of the *Alphaproteobacteria* class, more specifically the *Rhizobiaceae* family [1], and consists of rod-shaped, non-spore forming, gram-negative bacteria [2]. *Agrobacterium* is a very diverse genus comprising strains that can live in different environments, occurring mainly in soil and rhizosphere and even being capable of surviving under oligotrophic conditions such as pure water [3]. The diversity of this genus is also exemplified by the diverse effects it exerts on plants, with several *Agrobacterium* spp. being beneficial or non-pathogenic, while other species induce severe plant diseases, such as crown gall disease or hairy roots disease (HRD) on a wide range of host plants, often resulting in a considerable decrease in yield and associated economic losses [4–7].

*Agrobacterium* taxonomy has undergone many changes throughout the years. Initially, taxonomic classification of *Agrobacterium* members was based on the symptoms they caused on plant hosts. Bacteria capable of inducing tumours or crown galls were named *Bacterium tumefaciens* [8], those capable of inducing hairy roots were called *Phytomonas rhizogenes* [9], and non-pathogenic bacteria were assigned as *Bacillus radiobacter* [10]. Later, a new genus, *Agrobacterium*, was proposed by Conn [11] based on morphological and physiological similarities among these bacteria. This genus initially comprised the following three species: (i) *Agrobacterium tumefaciens*, causing tumours in a wide range of host plants; (ii) *Agrobacterium rhizogenes*, causing hairy roots in host plants; and (iii) *Agrobacterium radiobacter*, being non-pathogenic strains [9–11]. At that time, pathogenicity was considered the single trait to assign strains to specific *Agrobacterium* species [12]. Later on, three new species were added to the genus: *Agrobacterium larrymoorei*, which is capable of infecting figs (*Ficus* spp.) [13, 14]; *Agrobacterium rubi*, capable of infecting berries (*Rubus* spp.) [15, 16]; and *Agrobacterium vitis*, causing woody galls on grapevine (*Vitis* spp.) [17]. However, it was observed later on that pathogenicity was determined by the type of plasmid carried by the pathogenic strain, i.e., the tumor-inducing plasmid (pTi) for tumorigenic strains or the root-inducing plasmid (pRi) for rhizogenic strains [18, 19]. Moreover, the observation that tumorigenic agrobacteria could be ‘converted’ into rhizogenic agrobacteria by substituting one type of plasmid for another led to a revision of the classification system of *Agrobacterium* species [20]. All this indicated that a characteristic resulting from the arbitrary loss or acquisition of a plasmid was unstable and could not form the basis of formal taxonomic nomenclature.

Subsequently, mainly based on biochemical and serological tests, *Agrobacterium* species were classified into three biovars [21]. It should be noted that in the case of *Agrobacterium*, the term “biovar” does not have the usual meaning of a specific phenotypic form within a species, but rather corresponds to biological species within the genus [22]. Because the delineation in biovars is based mainly on the results of biochemical tests, tumorigenic, rhizogenic, and non-pathogenic strains could be assigned to the same biovar. With the dawn of the genomic era, Young et al. [12] proposed to merge the *Agrobacterium* genus with that of *Rhizobium* based on 16S rDNA (*rss*) analysis, since both formed a monophyletic clade, along with *Allorhizobium*. This reclassification was based on the analysis of *rrs* gene sequences and the lack of concordance between DNA-DNA hybridization (DDH), biochemical traits, and fatty acid profiles [12]. However, this suggestion was challenged by Farrand et al. [23] who claimed that only biovar 2 (*A. rhizogenes*) belongs to the genus *Rhizobium*, giving birth to the name of *Rhizobium rhizogenes*. The epithet “rhizogenes” did not imply pathogenic properties [1]. Eventually, biovar 3 was changed from *A. vitis* into *Allorhizobium vitis* [17, 24]. All these changes resulted in the fact that strains initially classified into three different biovars of the same genus, ended up being classified into three different genera: biovar 1 strains were classified as *A. tumefaciens*, biovar 2 strains as *R. rhizogenes*, and biovar 3 strains as *All*. *vitis* [25, 26]. This separation has been further confirmed by the absence of a linear chromid in *R. rhizogenes* and *All. vitis* strains, which is considered a critical taxonomic characteristic for *Agrobacterium* [27]. In contrast, *All. vitis* and *R. rhizogenes* strains carry a circular chromid (i.e., a replicating DNA molecule which has characteristic features of both chromosomes and plasmids) in addition to the chromosome [27, 28].

Considering the complexity of classification of *Agrobacterium* (and related genera) described above, there is clearly a need for tools to unequivocally assign strains to dedicated genera and species. And although DDH has been the standard to delineate species in the past, it has been shown that DDH often produces highly variable results [29]. Hence, it has been proposed that in addition to DDH and morphological, physiological, and biochemical properties, modern bacterial taxonomy should employ genetic information as the ultimate criterion to validly delineate bona fide species [30, 31]. Deeper taxonomic analyses, including the analysis of specific marker genes such as genes encoding the 16S rRNA and a DNA recombinase (*recA*), revealed that former *Agrobacterium* biovar 1 is not a homogeneous species but a complex of several genomic species, also called genomovars or genomospecies [32]. For this reason, it has been proposed to rename *Agrobacterium* biovar 1 to ‘*A. tumefaciens* species complex’ (Atsc) until these genomospecies are unequivocally named and delineated [33]. To date, the Atsc contains 15 genomospecies (Table 1), referred to as G1 to G9, G11, G13 to G15, G19, and G20 [34, 35]. Until now, three genomospecies have not received yet binomial name (G5, G6, and G13) [1, 35, 36), while for 12 genomospecies either a valid (formal) name has been assigned, or a species name has been suggested (Table 1). In order to be validated, suggested names need to be approved and published in one of the Validation Lists regularly compiled by the International Journal of Systematic and Evolutionary Microbiology (IJSEM) [37]. Additionally, new *Agrobacterium* species have been described with the help of whole genome sequencing (WGS) and directly received a binomial name without being assigned a genomospecies designation, including *Agrobacterium arsenijevicii* [1], *Agrobacterium rosae*, *A. nepotum*, *Agrobacterium bohemicum*, *Agrobacterium cavarae* [38, 39], and *Agrobacterium skierniewecense* [40].

**Table 1.** List of species originally classified as *Agrobacterium*, of which some were re-classified later (adapted from [1])

| Species<sup>a</sup> | GS<sup>b</sup> | Pathogenicity<sup>c</sup> | Reference strain<sup>d</sup> | Original host | Previous naming | References |
| --- | --- | --- | --- | --- | --- | --- |
| *Agrobacterium tumefaciens* | G1 | NP | ATCC 4720<sup>T</sup> | Soybean (*Glycine max*) and common bean (*Phaseolus vulgaris*) | *Agrobacterium fabacearum* | [39,41, 42] |
| *Agrobacterium pusense* | G2 | T | NRCPB10<sup>T</sup> | Chickpea (*Cicer arietinum*), Lawson cypress (*Chamaecyparis lawsoniana*) | *Rhizobium pusense* | [43, 44-47] |
| *Agrobacterium tomkonis* | G3 | CI | IIF1SW-B1<sup>T</sup> | Human patients |  | [35,48] |
| *Agrobacterium radiobacter* | G4 | NP | LMG 140<sup>T</sup> |  | *Rhizobium radiobacter*, *Agrobacterium tumefaciens* | [22, 33, 49] |
| NA | G5 | ND | CFBP 6626 |  |  | [43] |
| NA | G6 | ND | NCPPB 925 |  |  | [43] |
| “*Agrobacterium deltaense*” | G7 | ND | YIC 4121<sup>T</sup> | *Sesbania cannabina* |  | [50] |
| “*Agrobacterium fabrum*” | G8 | T | C58<sup>T</sup> |  | *Agrobacterium tumefaciens* | [51] |
| *Agrobacterium salinitolerans* | G9 | ND | YIC 5082<sup>T</sup> | *Sesbania cannabina* |  | [52] |
| *Rhizobium rhizogenes* | G10 | HR | ATCC 11325<sup>T</sup> | Apple (*Malus* sp.) | *Agrobacterium rhizogenes* | [9, 11] |
| *Agrobacterium rubi* | G11 | T | TR3<sup>T</sup> (NBR C 13261<sup>T</sup>) | Blackberry (*Rubus* sp.) |  | [14, 15] |
| NA | G13 | ND | CFBP 6927 |  |  | [51] |
| *Agrobacterium nepotum* | G14 | T | 39/7<sup>T</sup> | *Prunus cerasifera* | *Rhizobium nepotum* | [40, 53] |
| “*Agrobacterium viscosum*” | G15 | ND | ATCC<br>31113<sup>T</sup> | *Rosa* spp. |  | [36] |
| *Agrobacterium burrii* | G19 | ND | Rnr<sup>T</sup> | *Rosa* spp. |  | [36, 54] |
| *Agrobacterium shirazense* | G20 | ND | OT33<sup>T</sup> | *Rosa* spp. |  | [36, 54] |
| *Agrobacterium cavarae* | NA | NP | RZME10<sup>T</sup> | Corn (*Zea mays*) |  | [38, 39] |
| *Agrobacterium larrymoorei* | NA | T | AF 3.10<sup>T</sup> | *Ficus benjamina* |  | [14] |
| *Rhizobium albertimagni* | NA | ND | AOL15<sup>T</sup> | Surface of aquatic<br>macrophyte<br>*Potamogeton pectinatus* | *Agrobacterium*<br>*albertimagni* | [55] |
| *Agrobacterium*<br>*skierniewecense* | NA | T | Ch11<sup>T</sup> | Chrysanthemum<br>(*Chrysanthemum* sp.) | *Rhizobium*<br>*skierniewecense* | [40] |
| *Agrobacterium arsenijevicii* | NA | T | KFB 330<sup>T</sup> | Raspberry<br>(*Rubus idaeus*) |  | [56] |
| “*Agrobacterium bohemicum*” | NA | NP | R90<sup>T</sup> | Poppy seed<br>(*Papaver somniferum*) |  | [57] |
| *Agrobacterium rosae* | NA | T | NCPBPB<br>1650<sup>T</sup> | Grapevine (*Vitis finifera*),<br>raspberry (*Rubus idaeus*)<br>and blueberry<br>(*Vaccinium* sp.) |  | [58] |
| *Agrobacterium cucumeris* | NA | HR | O132T | Cucumber (*Cucumis sativus*) |  | [59] |
| *Agrobacterium leguminum* | NA | NA | MOPV5 | *Phaseolus vulgaris* |  | [60] |
| *Agrobacterium divergens* | NA | NP | LMG 31531 | Soil |  | [61] |
| *Agrobacterium vaccinium* | NA | T | B7.6 | Blueberry<br>(*Vaccinium corymbosum*) |  | [62] |
| "*Agrobacterium oryzihabitans*" | NA | NA | M15 | Rice (*Oryza sativa*) | *Rhizobium oryzihabitans* | [63, 64] |
| *Allorhizobium vitis* | NA | T | K309<sup>T</sup> | Grapevine (*Vitis vinifera*) | *Agrobacterium vitis* | [17, 65] |

<sup>a</sup>Species names in quotation marks have not yet been included in the Approved List of Bacterial names
<sup>b</sup>GS: Genomospecies affiliation of species originally classified in the Atsc, according to the *recA* phylogeny as described by Costechareyre et al. [33]; NA: not assigned
<sup>c</sup>NP: Non-pathogenic, T: tumor-inducing strains, HR: hairy root-inducing strains, CI: clinical isolate, ND: not determined
<sup>d</sup>Official type strains are indicated with superscript T.

However, neither 16S rRNA nor *recA* marker gene phylogeny provide the resolution required to unequivocally classify strains into *Agrobacterium* genomospecies. To further scrutinize taxonomy in such complex genera, multi-locus sequence analysis (MLSA), comparative genome hybridization (CGH), or core-genome phylogeny have been used [33, 40,43,51, 66]. More and more, overall genome relatedness indices (OGRI) that use genome-wide comparisons are commonly used to delineate prokaryotic species, such as ANI using BLASTN (ANIb) or MUMmer (ANIm) algorithms, genome BLAST distance phylogeny (GBDP), or tetranucleotide frequencies (TETRA) [26, 67, 68, 69]. Average nucleotide identity (ANI) values calculated for a limited set of conserved core genes (referred to as ANIo) have also been proposed to replace traditional DDHs [70].

In conclusion, even though recently a set of guidelines have been proposed for the description of new genera and species of rhizobia and agrobacteria [71], they have not been applied on a large collection of *Agrobacterium* strains to confirm their accuracy. Moreover, the taxonomic position of rhizogenic strains has also been poorly investigated with only a few studies available. The main objectives of this study were therefore to: (i) gain more insight into the taxonomic classification of *Agrobacterium* strains; (ii) evaluate and compare the ability of different DNA-sequence based tools for the delineation of (genomo)species, in particular within the AtSC; and (iii) examine the positioning of rhizogenic agrobacteria within the *Rhizobiaceae* family. To achieve this, we compared 579 publicly available genome sequences by means of different genetic analyses, including 16S rRNA and *recA* marker gene phylogeny, MLSA (including the housekeeping genes 16S rRNA, *atpD* encoding for the ATP synthase B subunit*, gyrB* encoding for a DNA gyrase subunit*, recA*, and *rpoB* encoding for an RNA polymerase subunit), WGS phylogeny, and OGRI values, such as ANIb, AAI, and nonmetric multidimensional scaling (NMDS) analysis based on and codon usage. Additionally, to increase our understanding of phylogenetic relationship of rhizogenic *Agrobacterium* strains, we included a selection of 24 (in-house) sequenced strains that were isolated from hydroponic crops showing HRD symptoms.

## MATERIALS AND METHODS

### Genome sequencing and collection of publicly available genome sequences

A total of 579 quality-checked (based on NCBI metadata annotation for contaminated, unverified, or fragmented) genome sequences were downloaded from the NCBI database in September 2023 using the get_assemblies v.0.10.0 package (Davis, 2020) [72], including 412 strains classified as *Agrobacterium*, 8 as *Allorhizobium* (ex *A. vitis*), and 159 as *Rhizobium* (S1 Table). The main focus was on strains previously classified as *Agrobacterium* and/or identified as rhizogenic strains (classified as *Agrobacterium* spp. or *R. rhizogenes* strains). The binomial names associated with each assembly, as retrieved in the NCBI database, were respected. In addition, the genome sequence of type strains of several *Rhizobium* and *Agrobacterium* strains were included to validate the confidence of the phylogenetic trees (S1 Table).

This collection of genome assemblies included a representative set of 24 rhizogenic *Agrobacterium* strains previously isolated from HRD-infested greenhouses from different plant hosts (tomato, cucumber and melon) and countries across Europe, the United States, Canada and Japan. [73, 74] (S2 Table). For details regarding sequencing and assembly, we refer to Kim et al. [74]. Additionally, three *Ensifer adhaerens* assemblies (*Rhizobiaceae*, Hyphomicrobiales, Alphaproteobacteria) were used as an outgroup for the downstream analyses (S1 Table). As a result, the complete set of genome assemblies used in downstream analysis was composed of 582 strains.

### Construction of phylogenetic trees based on 16S rRNA gene, *recA* gene, multilocus sequence analysis (MLSA), and whole genome sequences

Two phylogenetic trees were constructed based either on the single marker gene 16S rRNA (1,515 bp) or *recA* (1,092 bp). In addition, an MLSA-based tree was constructed based on a concatemer of the 16S rRNA gene and four housekeeping genes, i.e., *atpD* (1,485 bp), *gyrB* (2,442 bp), *recA* (1,092 bp), and *rpoB* (4,149 bp), yielding a final concatemer of 10,683 bp. These trees were built using the autoMLSA2 workflow developed by Davis et al. [75]. Briefly, the DNA sequence of the individual marker genes of reference strain *Agrobacterium* NCPP2659 were downloaded from the NCBI GenBank database and used as query to retrieve homologous genes from the genome assemblies of the other 581 strains used in this study (S2 Table). For the individual marker genes as well as for the concatemer, the sequences were aligned using MAFFT v.7 [76] with default parameters, and a phylogenetic tree was generated using IQTREE v.2.2 [77–79]. The -program parameter was set to “BLASTN”. When more than one copy of the query gene 16S rRNA was found, only the highest scoring BLASTN hit was retained. The phylogenetic tree was constructed applying the Maximum Likelihood (ML) method and was based on the most optimal model found and a bootstrap of 1,000 replicates.

Whole genome-based phylogeny was constructed using the Phylophlan v.3.0 package [80]. First, a database of UniRef90 core proteins for *Agrobacterium* was generated by setting the ‘g’ parameter into s Agrobacterium_tumefaciens, which resulted in 1,524 markers. Next, a configuration file was generated indicating that the mapping of the database markers to the input genomes should be conducted using DIAMOND [81], the multiple sequence alignment should be constructed with MAFFT, that sequences should be trimmed using TrimAl, and that the trees should be first generated using FastTree and then enhanced using RaxML. Subsequently, the phylogeny was built by indicating the parameter –‘diversity’ as “low” since the genomes analysed were of closely related species. The resulting phylogenetic tree was recalculated using 1,000 bootstrap replicates. The resulting phylogeny was constructed using a ML method and the CAT General Time Reversible (GTRCAT) approximation was used as nucleotide substitution model. The obtained phylogenetic trees were annotated using the iTOL platform [82]. If available, genomospecies assignments were adopted from Costechareyre et al. [34] and Bosmans et al. [73].

Four phylogenetic signal indices (Blomberg’s *K*, Pagel’s *l*, Moran’s *I* and Abouheif’s *C*_mean_) were calculated in order to determine if variation in the genomic guanine-cytosine (GC) content among agrobacterial strains was linked to their phylogenetic affiliation. These phylogenetic signal analyses used as inputs: (i) the ML MLSA tree trimmed to include only *Agrobacterium* strains as well as the *Ensifer* outgroup; (ii) a table including all the GC content percentages, obtained from the assembly entries at the NCBI (Table S3). The phylosig() function of the R package “picante” v.1.8.2 was used to calculate Blomberg’s *K* and Pagel’s *l* [83], while the R packages phylobase v.0.8.10 [84] and abouheif.moran() function of the R package “adephylo” v.1.1-13 were used to calculate Moran’s *I* and Abouheif’s *C*_mean_ (method = “Abouheif” and “oriAbouheif”, respectively) [85], all considering 1000 simulations.

### Genome relatedness indices calculation

All genomes were compared in a pairwise manner to evaluate their phylogenetic relationship based on two OGRIs, ANIb and average amino acid identity (AAI). ANIb values were calculated using the average_nucleotide_identity.py script from pyani v.0.2.10 [86]. The threshold applied initially to delineate species was 95% based on López-Guerrero et al. [30]. This threshold was further fine-tuned in this study using the bactaxR package by identifying medoid genomes, i.e., genomes presenting the smallest average dissimilarity among themselves, using different thresholds [87]. AAI values were calculated using CompareM v.0.1.1 [88] under default settings using the aai_wf command, which uses protein coding sequences (CDS) predicted by Prodigal [89]. Then an all-*vs.*-all reciprocal sequence similarity search was conducted by DIAMOND [81] and finally pairwise AAI values based on the orthologous fraction of the two genomes was obtained. The threshold applied to delineate species in the AAI analysis was 95% based on Fan et al. [90]. Resulting distance matrices were visualized with heatmaps generated with the R package pheatmap v.1.0.12 in the RStudio v.3.3.0 platform [91–93].

### Analysis of codon usage patterns

Possible differences in the codon usage patterns among ANIb groups were visualized by principal component analysis (PCA), as implemented by the prcomp() function of the ‘stats’ R library [91]. The input data for the PCA was a data frame of the normalized relative synonymous codon usage (NRSCU) values calculated for the genomes included in this study using the Dynamic Codon Biaser webserver (http://www.cbdb.info/; [94]. PCA loading plots and scores plots were created using the ‘ggplot2’ R library [95].

## RESULTS

### Phylogenetic analysis of single marker genes

To have a comprehensive understanding of the phylogenetic relationships within the *Agrobacterium* genus, and the position of rhizogenic agrobacteria in the *Rhizobiaceae* family in particular, we analysed the genome sequence of 582 strains, including 24 in-house sequenced rhizogenic *Agrobacterium* strains. In a first approach, the phylogenetic relationships between strains, and more in particular, the association with previous genomospecies assignment was assessed based on 1,515 bp of the 16S rRNA gene (S1 Fig). The best substitution model for the 16S rRNA tree was Transition 3 model with empirical base frequencies and invariable site plus FreeRate with three categories model (TIM3+F+I+R3), based on Akaike information criterion (AIC) and Bayesian information criterion (BIC) scores of 17094.858 and 17158.736, respectively.

The 16S rRNA-based phylogeny showed that almost all strains clustered in three well-supported clades (>70% bootstrap values, 94.2% sequence similarity): (i) the R clade containing all the *Rhizobium* strains; (ii) a homogenous AV clade containing all the *All. vitis* (former *A. vitis*) strains; and (iii) the *Agrobacterium* clade containing all bona fide *Agrobacterium* strains. When zooming in on the R-clade, a subclade (R2) containing very closely related strains (>80% bootstrap values, 98.2% similarity) could be observed. This subclade contained all strains classified as *R. rhizogenes* (former *A. rhizogenes* or *Agrobacterium* biovar 2), as well as four *Agrobacterium* sp. strains. Interestingly, the R2 subclade also included *R. radiobacter* K84^T^, and the type strain of *R. rhizogenes* (i.e. strain NBRC 13257^T^ = LMG150^T^). In the *Agrobacterium* clade, seven well-supported subclades were identified (>80% bootstrap values, 96.8% similarity). The first cluster (Agr1_16S rRNA_) included strains that are part of the so-called *Agrobacterium rubi-larrymoorei* subclade, namely *A. vaccinii*, *A. rubi*, *A. skierniewicense*, *A. rosae*, and *A. bohemicum* strains as well as their respective type strains. The other clusters (Agr2-7_16S rRNA_) contained strains of different genomospecies (G1, G2, G3, G4, G5, G7, G9, G14, G19, and G20) that seemed to be interspersed throughout the clusters, except for Agr6_16S rRNA_ that was composed mainly of strains classified as *A. larrymoorei*.

Next, a phylogenetic tree was constructed based on a 1,092-bp alignment of the *recA* gene (S2 Fig). The best substitution model for the *recA* tree was Transition 3 model with empirical base frequencies and invariable site plus FreeRate with five categories model (TIM3+F+I+R5), based on AIC and BIC scores of 71080.772 and 71160.704. The *recA*-based phylogenetic tree presented two strongly supported clades (>80% bootstrap values, 91.5% similarity): (i) a clade containing most *Rhizobium* strains and (ii) the polyphyletic clade containing the *All. vitis* and *Agrobacterium* strains. Similar to the 16S rRNA tree, the *Rhizobium* clade could be divided into two subclades (R1 and R2), with subclade R2, mainly consisting of rhizogenic *Rhizobium* strains, including the type strain *R. rhizogenes* NBRC 13257^T^ (=LMG150^T^) and *R. radiobacter* K84. The *Allorhizobium*/*Agrobacterium* main clade could be further sub-divided into the *All. vitis* subclade (including the type strain *All. vitis* NCPPB 3554^T^) and the *Agrobacterium* subclade. The *Agrobacterium* subclade presented two clusters, being the *A. rubi*-*larrymoorei* cluster with nine groups and the Atsc cluster with 18 groups (≥84% similarity; >80% bootstrap values). The Atsc subclade was mainly composed of monophyletic groups consisting either of distinct genomospecies (based on the *recA* phylogeny reported by Costechareyre *et al.* [34]),. However, strains identified as “*A. deltaense*” and *A. salinitolerans* were dispersed into three (Atsc8-10, with Atsc10 containing the type strain “*A. deltaense*” YIC 4121^T^) and two (Atsc13-14, with Atsc13 containing the type strain *A. salinitolerans* YIC 5082^T^) polyphyletic groups, respectively. Similarly, *A. pusense* strains were split in two polyphyletic groups (Atsc15-16), with Atsc16 including the type strain LMG 25623^T^ (=NRCPB10^T^) and Atsc15 being composed of strains isolated from Pigeon Pea roots in India [96]. Additionally, Atsc11 contained strains with no previously assigned genomospecies affiliation nor any species names validly published or pending for approval nomenclature. Overall, the genomospecies delineation presented by the *recA* phylogeny indicated that this marker does not provide sufficient resolution to unequivocally classify all *Agrobacterium* strains in dedicated monophyletic genomospecies.

### Phylogenetic analysis using multiple marker genes using MLSA- and WGS-based trees

To overcome possible bias resulting from the use of single marker genes, an MLSA was conducted based on a concatemer sequence (10,683 bp in size) of five house-keeping genes (Fig 1). The best substitution models for the MLSA tree were Transition 3 model with empirical base frequencies and invariable site plus FreeRate with four categories model (TIM3+F+I+R) for 16S rRNA, Transition model 3 with empirical base frequencies and invariable sites plus FreeRate with six categories model (TIM3+F+I+R6) for *atpD* and *rpoB*, general time reversible model with unequal rates and unequal base frequencies and empirical base frequencies with invariable sites plus FreeRate with five categories model (GTR+F+I+R5) for *gyrB*, and Transition model 3 with unequal and empirical base frequencies with invariable sites plus FreeRate with five categories model (TIM3+F+I+R5) for *recA.* These models were selected based on BIC scores of 19104.979, 291897.533, 197574.819, and 69398.569 and AIC scores of 19030.454, 291778.075, 197470.409, and 69318.637 for 16S rRNA, *atpD* and *rpoB*, *gyrB*, and *recA*, respectively.

**Figure 1.**
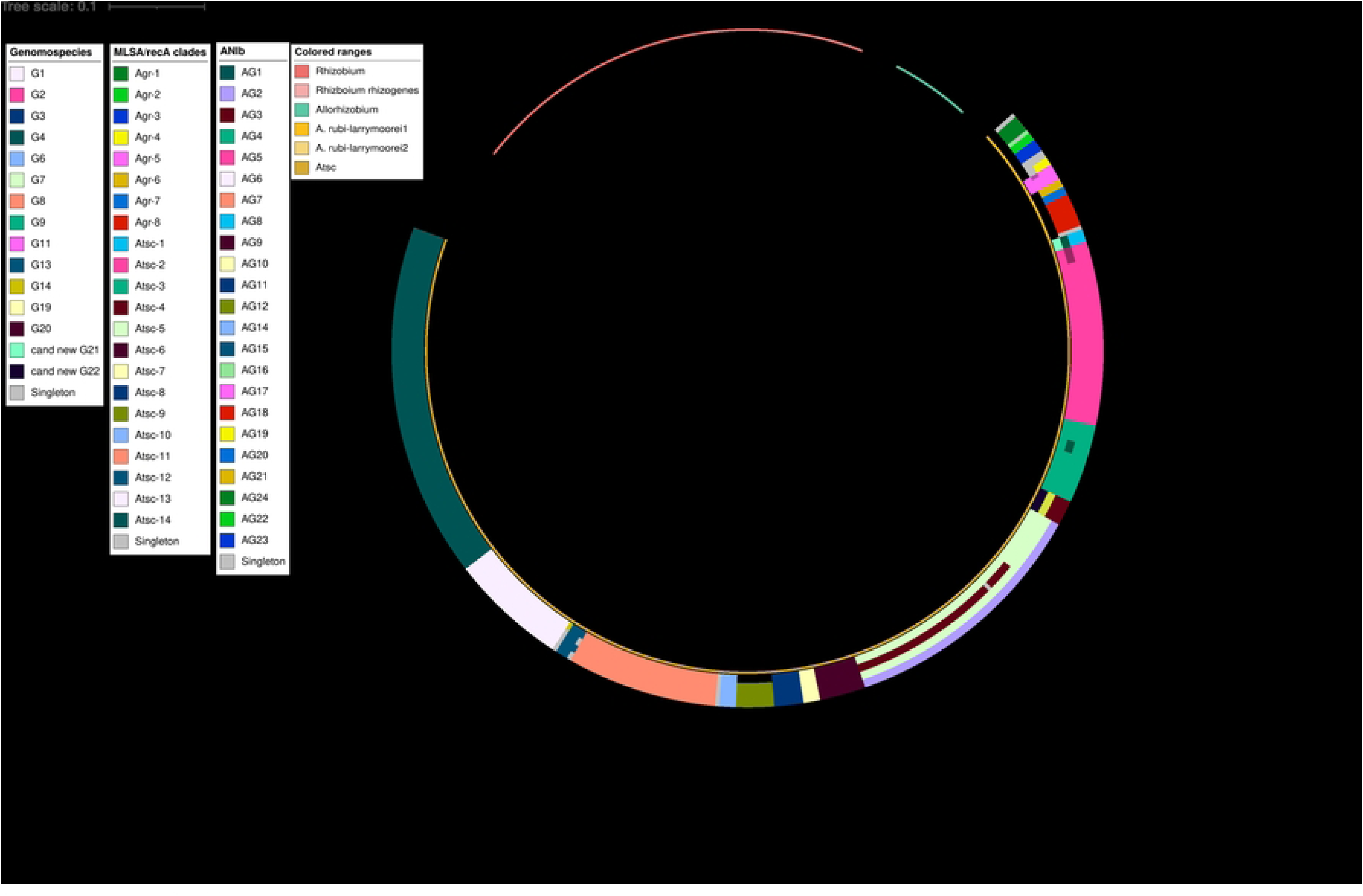
Maximum likelihood (ML) tree based on a multilocus sequence analysis of *Agrobacterium*, *Allorhizobium*, and *Rhizobium* strains. A ML phylogeny of 582 strains including 412 *Agrobacterium*, 8 *Allorhizobium*, and 159 *Rhizobium* strains is shown. The phylogeny was built from the concatenated alignment (10,683 bp) of five chromosomal genes (16S rRNA, *atpD*, *gyrB*, *recA*, and *rpoB*) using IQTREE. Bootstrap support value ranging from 70% to 100% based on 1,000 repetitions is represented by thicker branches. Leaves have been highlighted according to genus affiliation. Leaves have also been annotated in the outer rings according to (from inside to outside) original genomospecies affiliation, MLSA subclades delineation, and ANIb subgroups (using a 94.0% identity threshold). Type strains are highlighted in light-blue background and bold font (see detailed annotation in online version https://itol.embl.de/shared/pme_bim). The scale bar represents the number of expected substitutions per site under the best-fitting model for each gene (TIM3+F+I+R for 16S rRNA, TIM3+F+I+R6 for *atpD* and *rpoB*, GTR+F+I+R5 for *gyrB*, and TIM3+F+I+R5 for *recA*; see details in Materials and Methods). Strains that did not cluster with any other are annotated as STN for singleton. *E. adhaerens* strains were used as outgroup.

In general, the MLSA phylogeny showed a similar structure as the *recA* tree, and presented two strongly supported clades (>80% bootstrap values, ≥91% similarity): (i) the *Rhizobium* clade, which could be subdivided in a subclade R2 containing closely related (mostly *R. rhizogenes*) strains and another subclade, consisting of more distantly related *Rhizobium* strains; and (ii) the clade containing all the *All. Vitis* and *Agrobacterium* strains. Similar to the *recA* phylogeny, the *All. vitis*/*Agrobacterium* clade could be further subdivided into the *All. Vitis* subclade (including the type strain NCPPB 3354^T^), the *Agrobacterium* subclade, and a set of closely related recently described genera including *Heterorhizobium*, *Paenirhizobium*, *Peteryoungia*, *Affinirhizobium*, *Neorhizobium*, and *Alirhizobium* [64].

When looking deeper into the *Agrobacterium* clade, eight clusters were part of the *A. rubi*-*larrymoorei* subclade (Agr1-8) and contained bona fide species, while 14 clusters were grouped together in the Atsc subclade and included mainly known GSs, bona fide validly published species names or species names awaiting for nomenclature. Similarly, strains identified as *A. pusense*, *A. salinitolerans*, and “*A. deltaense*” also clustered monophyletically. Noteworthy, strains classified as *A. leguminum* clustered together with “*A. deltaense*” strains. Additionally, clades Atsc1 and Atsc4 contained strains with no previous GS affiliation clustered apart.

To resolve the ambiguities between the *recA* and MLSA trees, a WGS phylogeny was constructed. The phylogeny reconstruction was based on a concatemer of 852,736 bp that presented 851,130 informative sites. The likelihood value of the final tree obtained was –74728904.543924. The WGS phylogeny presented three strongly supported clades (>80% bootstrap values, 92% similarity): (i) the *Rhizobium* clade; (ii) the *Allorhizobium* clade; and (iii) the *Agrobacterium* clade. Similar to the MLSA phylogeny, *Heterorhizobium*, *Paenirhizobium*, and *Peteryoungia* clustered closely with *Allorhizobium*. The WGS phylogeny showed an easily distinguishable pattern in the *Agrobacterium* clade and displayed nine and 15 groups in the *A. rubi-larrymoorei* and the Atsc subclades, respectively (Fig 2). Most of the defined subclades coincided with genomospecies assignment, validly published species or species waiting for name validation. Similar to the *recA* and MLSA phylogenies, strains identified as “*A. deltaense*” and *A. leguminum* clustered together in Atsc14_WGS_. Noteworthy groups Agr9, Atsc7, Atsc10, and Atsc12 contained strains with no previous GS assignation.

**Figure 2.**
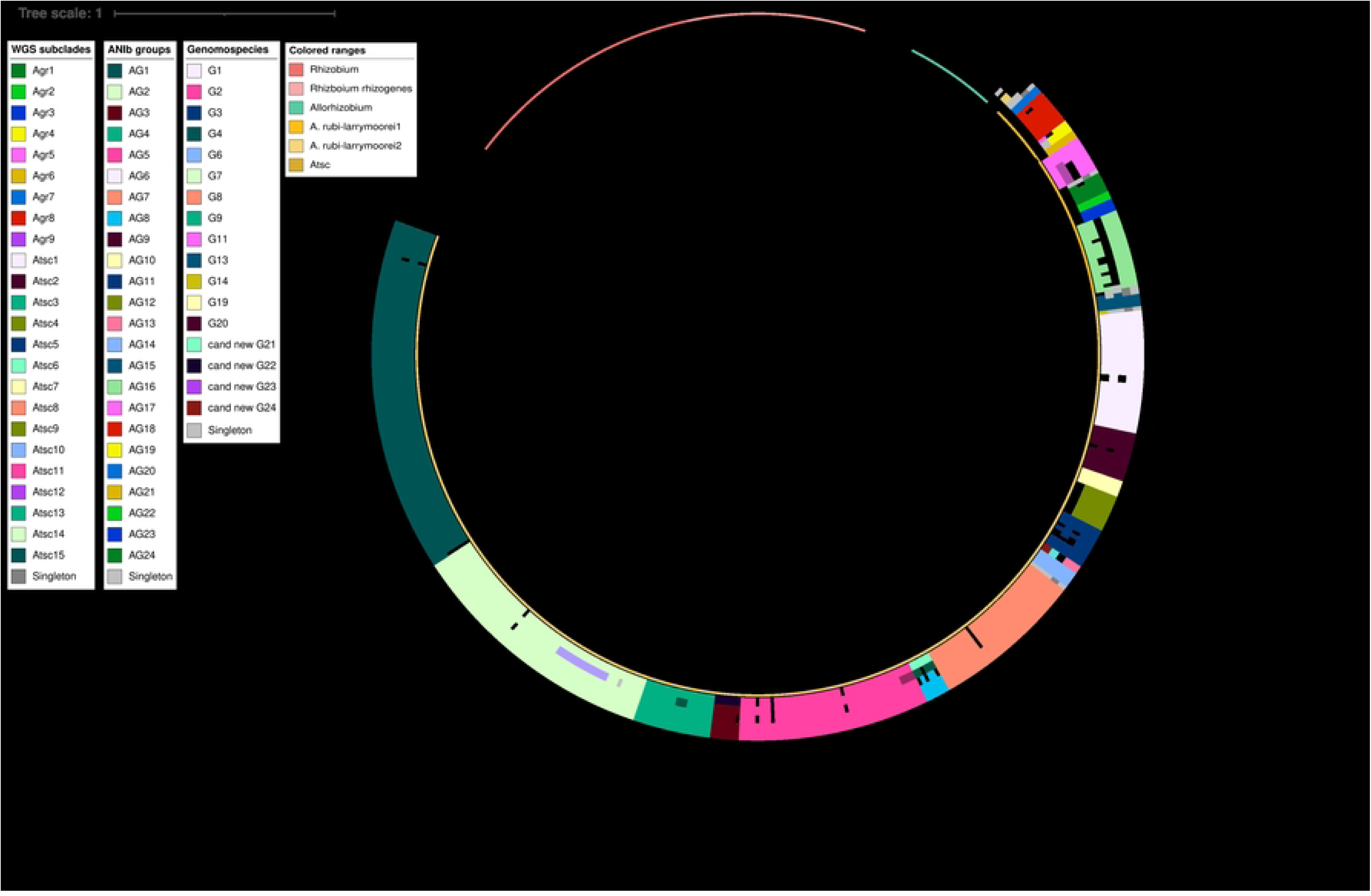
Whole-genome sequence (WGS) phylogenetic tree of *Agrobacterium*, *Allorhizobium*, and *Rhizobium* strains. The tree was obtained using the Maximum likelihood (ML) method. The phylogeny was reconstructed initially with FastTRee and then enhanced with RAxML applying the ML method and a CAT General Time Reversible (GTRCAT) approximation. The phylogenetic analysis was based on a concatenated alignment of the most discriminative amino acid positions of 1,524 markers conserved among all 582 sequenced genomes (412 *Agrobacterium*, 8 *Allorhizobium*, and 159 *Rhizobium* and three *E. adhaerens* strains used as outgroup) identified with Phylophlan 2. Bootstrap support values ranging from 70% to 100% based on 1,000 repetitions is represented by thicker branches. Leaves have been highlighted according to genus affiliation. Type strains are highlighted in a light-blue background and bold font (see detailed annotation in online version https://itol.embl.de/shared/pme_bim). Leaves have also been annotated in the outer rings according to (from inside to outside) genomospecies affiliation, WGS subclades delineation, and ANIb subgroups. The scale bar represents the number of expected substitutions per site.

**Figure 3.**
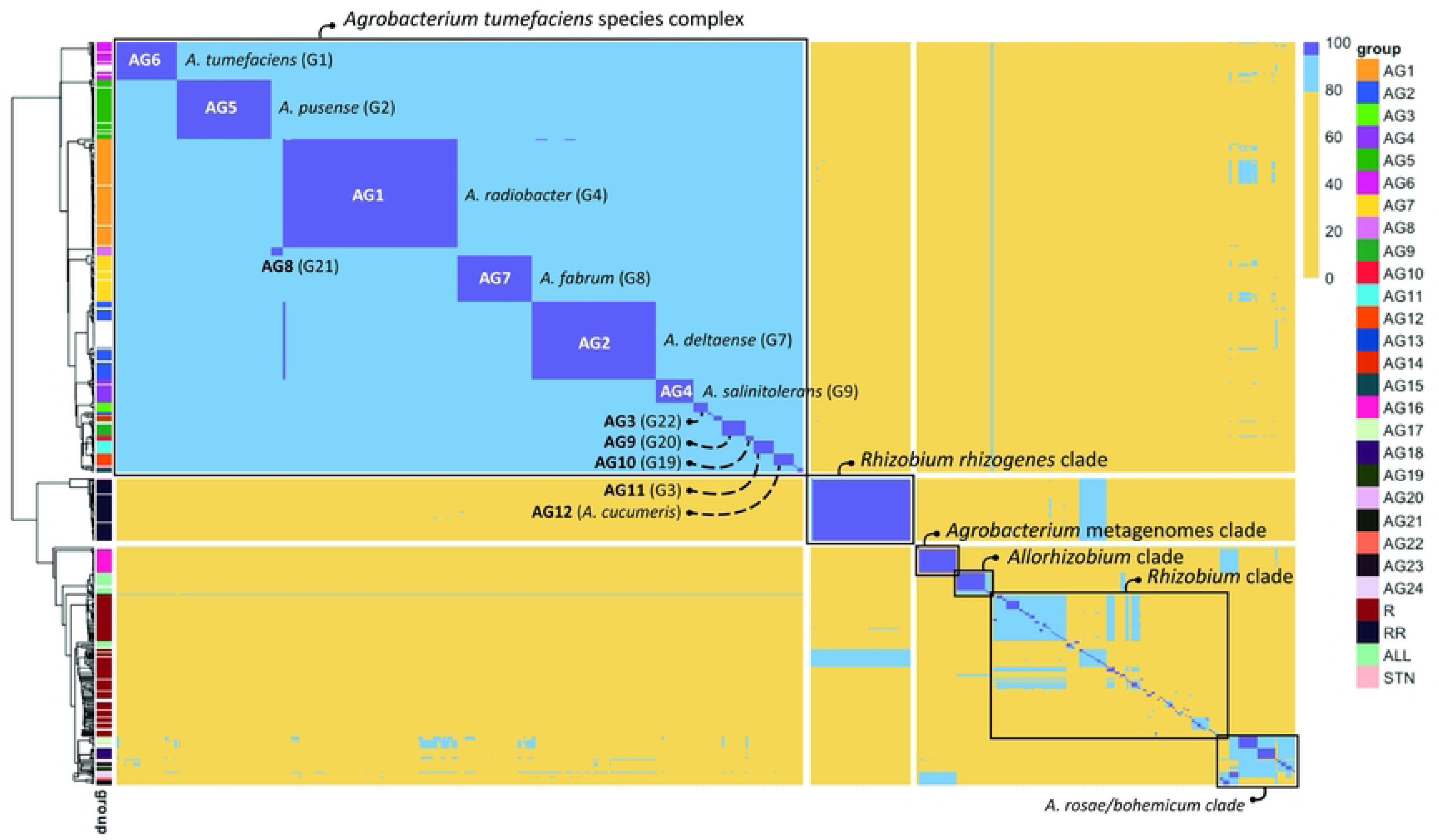
Heat map of average nucleotide identity using BLAST (ANIb) pairwise comparison. Pairwise ANIb values are shown of 412 *Agrobacterium*, 8 *Allorhizobium*, and 159 *Rhizobium* genomes. A total of 24 ANIb groups (AG1-24) were observed in the *Agrobacterium* genus. In addition 7 strains formed singletons, i.e. did not show an average nucleotide identity above 94% with any other strain. Pairwise comparison values ≥ 94.0% are highlighted in dark blue(corresponding to species level) and those between 80.0% and 94.0% are highlighted in light blue(corresponding to genus delineation). The hierarchical tree was constructed based on the pairwise comparison values using a “complete” clustering algorithm. The hierarchical tree has been annotated with ANIb groups based on pairwise comparison values of ≥ 94.0% among strains.

### Assessment of phylogenetic signals based on GC-content, presence/absence of *Agrobacterium* genes, and codon usage bias analysis

The phylogenetic signal analysis involving GC content of all the *Agrobacterium* strains from the MLSA phylogeny yielded statistically significant results in all four metrics, more specifically Blomberg’s K=2.6·10^-6^ (p-value=0.001), Pagel’s Lambda=0.976 (p-value=2.73·10^-202^), Moran’s *I*=0.81 (p-value ≤ 0.001) and Abouheif’s *C_mean_*=0.849 (p-value ≤ 0.001), meaning that the distribution of GC values across the MLSA phylogeny was not random but, instead, depended on the phylogenetic relatedness between strains (i.e., close relatives displayed more similar values among themselves than with unrelated strains).

Finally, it was also assessed whether the codon usage of *Agrobacterium* strains could be correlated to genomospecies classification (S4 Fig). Inspection of the results revealed a wide variation of the PCA scores within most ANIb groups, which resulted in the presence of a number of outliers and precluded a clear separation of most groups. Therefore, it was concluded that codon usage patterns might not be a good criterion for taxonomic delimitation.

### Use of overall Genome Relatedness Indices for species delineation in the *Agrobacterium tumefaciens* species complex

In order to better determine the species boundaries within the *Agrobacterium* genus, an ANIb was conducted as a complementary method since it is considered the golden standard for prokaryotic species definition [32]. The strains examined in this study yielded 160,801 pairwise comparisons showed pairwise distance values ranging from 68.06%% to 99.99% (S4 Table). At first glance and in contrast with the *recA*, MLSA, and WGS phylogenies, the ANIb-based hierarchical tree suggested that *All. vitis* and *Rhizobium* strains seemed to be more closely related to each other and more distant from *Agrobacterium* (Fig 4). Remarkably, *Agrobacterium* species composing the *A. rubi*-*larrymoorei* clade of the three phylogenies (*recA*, MLSA, WGS) reconstructed, clustered within the *Rhizobium* clade in the ANIb-based tree.

Several studies have indicated that an ANIb cut-off value of 94-96% similarity should be applied to determine a species threshold among prokaryotes [30, 32]. To establish a more accurate cut-off value for *Agrobacterium* species delineation, we analysed the ANIb-values of all strains and observed a clear gap in the distribution of ANIb values between 93.7% and 94.2% (S3 Fig). For this reason, we have set the cut-off to delineate distinct subclades at an ANIb of 94.0%.

Using the optimized ANIb cut-off for species delineation, the *Agrobacterium* clade presented 24 groups (AG1-24) and 7 singletons, with pairwise distance values ranging from 76.58% to 99.99%. Annotating the ANIb results on the *recA*, MLSA, and WGS phylogenies indicated that MLSA and WGS analysis resulted in a more accurate delineation in species compared to *recA* phylogeny. In the WGS phylogeny, ANIb annotation helped to better delineate clades, mainly merge and form monophyletic groups. For example, ANIb allowed: (i) merging Agr3_WGS_ strains with neighbouring singletons and form a single group of *A. vaccinii* strains; (ii) merging all neighbouring *A. rubi* strains into a single group Agr5_WGS_; (iii) merging all neighbouring *A. pusense* strains into a single group Atsc11_WGS_; (iv) merging all neighbouring *A. salinitoelrans* strains into a single group Atsc13_WGS_; and (v) merging all neighbouring *A. radiobacter* strains into a single group Atsc15_WGS_. Strains in the *All. vitis* clade showed an identity range of 77.25%-99.99% and revealed three subclades and three singletons at a 94% identity threshold (Fig 4). The groups as defined based on ANIb analysis corresponded perfectly with the MLSA and WGS phylogeny.

In addition, pairwise AAI values were calculated to evaluate the capacity of this OGRI to delineate species in the Atsc. AAI analysis assessed the homology (evaluating both sequence similarity and alignment length) of shared CDSs. The strains examined in this study showed pairwise AAI distance values ranging from 67.3% to 99.99% (S4Table). Applying a cut-off of 80% similarity for genus delineation as advised by Liu et al. [97], 13 distinct clusters could be identified: (i) a homogenous cluster containing all *All. vitis* strains; (ii) a cluster containing strains of the *A. rubi*-*larrymoorei* clade of the three phylogenies reconstructed; (iii) three remaining *Agrobacterium* clusters, one of them composed of only two strains (Azo12 and AGB0); and (iv) nine *Rhizobium* clusters, one of them being composed exclusively of strains identified as *R. rhizogenes*. By applying a threshold identity value of 95% to delineate species, as recommended by Fan et al. [90], a total of 27 clusters (9 in the *Rhizobium* clade, 2 in the *Allorhizobium* clade, and 16 in the Atsc clade) and 55 singletons were identified (S5 Table). Remarkably, one singleton identified as *Agrobacterium* sp. a22-2 and two clusters with strains identified as *Agrobacterium* (*Agrobacterium* sp. MA01 and RAC06, and *A. albertimagni* AOL15 and *Agrobacterium* sp. SCN 61-19) were observed closely related to other *Rhizobium* singletons.

## DISCUSSION

The main objective of this study was to further elucidate and delineate (genomo)species in the *Agrobacterium* genus and to have better insight into the taxonomic positioning of rhizogenic strains, i.e. strains that induce HRD. To achieve this, a large set of more than 400 publicly available *Agrobacterium* genomes originally identified as members of genus *Agrobacterium* were analysed using several analysis tools that are frequently used to infer phylogenetic relationships, which also allowed evaluating their usefulness for species delineationwithin *Agrobacterium*. Because we were particularly interested in the taxonomic positioning of rhizogenic strains (i.e., strains causing HRD), we also included a set of 63 *R. rhizogenes* strains and 24 in-house sequenced rhizogenic *Agrobacterium* strains (previously classified as *A. rhizogenes*). In order to position rhizogenic *Agrobacterium* and *Rhizobium* strains in a broader phylogenetic context, we also included 18 *All. vitis* (former *A. vitis*) strains, and 96 *Rhizobium* type strains in this study. The phylogenetic relationship among *Agrobacterium*, *Allorhizobium*, and *Rhizobium* species was evaluated by generating four phylogenetic trees: two based on single marker genes (16S rRNA or *recA* gene), one comprising concatemers of the 16S rRNA gene and four housekeeping genes (*rpoB*, *recA*, *gyrB*, and *atpD*), and one using WGS data. Additionally, the pairwise ANI and AAI were calculated for all strains, and among *Agrobacterium* strains it was assessed whether presence/absence of genes, codon usage, or GC-percentage contain a phylogenetic signal. The results obtained have confirmed previously reported findings, but also provided new insights in the positioning of rhizogenic agrobacteria within the taxonomy of the Atsc. This study also allowed to evaluate the accuracy of different analyses to infer taxonomy within the genus *Agrobacterium*.

### Single-marker phylogeny does not provide sufficient taxonomic resolution in the *Agrobacterium* genus

The sequence of the 16S rRNA gene was used to infer taxonomic classification since it is a well-known and valuable marker for bacterial taxonomy [98]. This marker is commonly included in polyphasic taxonomy approaches for bacteria due to the availability of universal primers and inherent high copy number in bacterial genomes [99]. In our study, the 16S rRNA-based phylogeny proved useful to distinguish strains at the genus level, and presented three large clades corresponding to *Rhizobium*, *Agrobacterium*, and *Allorhizobium*. Interestingly, a few strains identified as *Rhizobium* did not cluster in any of these large clades, indicating that these strains do not belong to genus *Rhizobium*, refuting their original classification. These strains have been recently moved into new genera including, (i) *R. subbaraonis* JC85^T^ and *R. rhizolycopersici* DBTS2^T^ to *Mycoplana*; (ii) *R. album* NS-104^T^ to *Metarhizobium* (also considered at basal position in *Rhizobium* phylogeny); (iii) *R. wenxinae* DSM 100734^T^ and CGMMCC 1.15279 as well as *Agrobacterium* sp. Ap1 to *Alirhizobium*; and (iv) *All. pseudoryzae* DSM 19479^T^ and *R. rhizoryzae* DSM 29514^T^ to *Affinirhizobium*. Nevertheless, it is a well-known fact that 16S rRNA does not provide sufficient resolution for species delineation in these complex genera [25, 100]. This is exemplified by some type strains of different *Agrobacterium* genomospecies sharing a nearly identical 16S rRNA sequence. For example, the type strains *A. burrii* Rnr^T^ (G19) and *A. tumefaciens* LMG 140^T^ (now *A. radiobacter*, G4) shared 99.9% 16S rRNA gene similarity, which is higher than the recommended 98.7% threshold to delineate (genomo)species [38]. This poor capacity to delineate prokaryotic species might be driven by the nucleotide substitution rates for the 16S rRNA gene that vary among different bacterial groups [25]. In the case of the *Rhizobiaceae* family, the gene substitution rates are low, meaning that this marker is often too conserved, resulting in few variant positions within a genus or species [25]. Moreover, it has been reported that certain segments of the 16S rRNA gene among *Rhizobiaceae* species may have a history of lateral transfer or recombination, thus presenting a mosaic structure that blurs accurate species delineation using this marker [101].

In general, protein-encoding genes are considered more informative for taxonomic classification [25, 102]. Aiming for a clearer delineation of *Agrobacterium* (genomo)species, a phylogenetic analysis was conducted using the housekeeping gene *recA*. This gene encodes a DNA recombinase and was initially used for assigning strains in the Atsc to a specific genomospecies [25, 33]. Our *recA* phylogeny was mainly congruent with that presented by Costechareyre et al. [33], who defined thirteen distinct clades, nine of which were identified as *Agrobacterium* genomospecies G1-9 and four that were identified as *A. larrymoorei*, *A. rubi*, *A. rosae*, and *A. vitis* (later renamed to *All. vitis*). Even though the use of *recA* was initially used to differentiate between genomospecies, our results showed that *recA* was not always sufficient to make a clear delineation between genomospecies. For instance, in our study the *recA*-based phylogenetic tree did not generate monophyletic groups for some (genomo)species including *A. vaccinii*, *A. rubi* (G13), *A. salinitolerans* (G9), *A. pusense* (G2) and “*A. deltaense*” (G7). For that reason, we conducted an (*in silico*) MLSA that concatenated the 16S rRNA gene and four housekeeping genes (*rpoB*, *recA*, *gyrB,* and *atpD*). The application of multiple genes to resolve taxonomic issues has been previously documented to overcome the conflicting branching patterns of individual genes and has even been recommended for the genus *Agrobacterium* [40]. Although the MLSA tree was mainly congruent with the *recA* tree, it was clear that MLSA provided a higher taxonomic resolution. For instance, the MLSA tree clustered together all strains identified as *A. pusense* (G2) and those identified as “*A. deltaense*” (G7), unlike the *recA* phylogeny in which they were split in two clusters. Interestingly, strains identified as *A. leguminum* (including type strain *A. leguminum* MOPV5^T^) were part of this G7-containing MLSA cluster.

Considering that the MLSA tree topology as well as genomospecies delineation was largely supported by both the WGS tree and ANIb analysis, it can be concluded that an MLSA-based tree outperforms phylogenetic trees based on either 16S rRNA or the *recA* genes, which are clearly not sufficient for accurate taxonomic positioning of strains within the *Agrobacterium* genus.

### A combination of phylogenetic analyses provides support to “new candidate genomospecies” and allows identification of misclassified strains

The MLSA tree identified strains and subclades that clustered separately from previously reported genomospecies, suggesting the identification of new and previously unreported genomospecies. We provided support for this hypothesis by using a combination of ANIb analysis and WGS phylogeny. Our results indicated that, in the *Agrobacterium* clade of the WGS tree, strains *Agrobacterium* sp. DE0009, *A. tumefaciens* UBA5877, *A. tumefaciens* UBA6714, *A. tumefaciens* 6N2, and *A. tumefaciens* KCFJK1736 were clustered in subclade Atsc10_WGS_. This clustering was supported by ANIb, since all the strains in question were part of the ANI-group AG8 at the cut-off (94.0%) defined in this study. For these reasons, we suggest renaming it as new candidate genomospecies G21.

Similarly, strains in both Atsc4_MLSA_ and Atsc12_WGS_ group also clustered together in the same ANIb AG3 group, indicating that these strains belong to the same species. These strains were isolated from hospital intensive care unit sinks in the United States of America in 2023. As a result, we suggest considering these strains as new candidate genomospecies G22.

Moreover, a cluster composed exclusively of MAGs strains was observed in the *A. rubi-larrymoorei*. The ANIb results also pointed out that these strains are part of the same AG16 group. Based on these findings, we suggest considering these strains as new candidate genomospecies G23, supported by Minute 10 of the 2022 meeting of the Subcommittee on the Taxonomy of Rhizobia and Agrobacteria, wherein the SeqCode was endorsed [103]). Additionally, strains *A. tumefaciens* BIN17 and *Agrobacterium* sp. SOY 23 clustered apart in the *recA* (Atsc11*_recA_*) and WGS (Atsc7_WGS_) phylogenies as well as in the ANIb AG13 group. These results suggests that these strains could be also part of a new candidate genomospecies G24. So far, it seems *Agrobacterium* BIN17 and SOY23 are non-pathogenic species since they were isolated from a lignocellulose-decomposing bacterial consortium from soil associated with dry sugarcane straw and soybean rhizosphere soil [104].Additionally, strain ST15.13.057 is hypothesized to be a new genomospecies since it did not cluster with any other strain in any of the phylogenies reconstructed nor in the ANIb analysis. This strain was isolated in a HRD-infested tomato greenhouse. Interestingly, compared to several other rhizogenic *Agrobacterium* strains, ST15.13.057 seems to be a highly aggressive strain resulting in severe symptoms of excessive root growth (Kim et al., unpublished data). It is also characterized by a comparatively large genome size (Supplementary Table S1). Nevertheless, more related strains are needed in order to propose ST15.13.057 as a new genomospecies according to the rules provided by the International Committee on Systematics of Prokaryotes (ICSP) Subcommittee on the Taxonomy of Rhizobia and Agrobacteria [105].

Although several complementary analyses (including MLSA- and WGS-based phylogeny, and ANIb analysis, confirm our hypothesis that strains mentioned above are members of new genomospecies, additional analyses (including fatty acid profiles, cell wall composition, and exopolysaccharides, as well as morphological, biochemical, and enzymological characterization) are required to confirm that there is sufficient difference among the strains of these clades to formally describe them as new species within the genus *Agrobacterium* [106].

Finally, our analysis also provides support for recently described new species. Indeed, the MLSA-and WGS-based phylogeny showed that strains previously assigned to G3 were clearly divided in two separate subclades. While strains of subclade Atsc8_MLSA_ or Atsc5_WGS_ were assigned to *A. tomkonis* (G3), strains of the other subclade were reclassified into *A. cucumeris* [59]. This was also supported by the ANIb analysis that showed <94.0% identity among strains of both subclades.

In addition to identifying candidate new genomospecies, MLSA analysis enabled a more accurate classification into genomospecies than *recA* analysis. For instance, while *recA* phylogeny positioned some outside the *A. pusense* (G2) clade (Atsc16*_recA_*), the MLSA phylogeny positioned these strains within the G2-containing subclade (Atsc2_MLSA_). Classification of these strains as G2 was also supported by ANIb analysis and WGS phylogeny. A similar situation was observed with “*A. deltaense*” (G7) strains that were separated into two clusters in the *recA* phylogeny (Atsc8*_recA_*and Atsc10*_recA_*) but were merged in the MLSA phylogeny (Atsc5_MLSA_).

Within the *Agrobacterium* clade, a set of strains seemed to be misclassified. Such was the case for two *A. larrymoorei* strains, SORGH AS 1126 and SORGH AS974, that actually clustered apart in both the MLSA and WGS phylogenies and showed <94% nucleotide identity (namely 78.3-78.6%) with other *A. larrymoorei* strains (including the type strain). Likewise, strains previously identified as *A. leguminum*, including the type strain *A. leguminum* MOPV5^T^, clustered together with “*A. deltaense*” (G7) strains in both the MLSA and the WGS phylogenies. This suggests that *A. leguminum* and *A. deltaense* strains both belong to the same species (previously assigned as G7), which is also supported with ANIb values >95% between *A. deltaense* and *A. leguminum* strains. Similarly, other strain identifiers would need to be adjusted, including *R. rhizogenes* strains B 4.1 and SBV 302 82 into *A. tumefaciens* B 4.1 and *A. burrii* SBV 302 78 2, and *R. oryzihabitans* M15 should be *A. oryzihabitans* (G13) [64]. Likewise, all ST in-house strains need to be reclassified according to their respective *Agrobacterium* cluster. Also, outside the *Agrobacterium* clade, our study identified strains that seemed to be misclassified. For instance, *A. albertimagni* AOL51^T^, which clustered closely with the *Agrobacterium* clade in the 16S rRNA-based phylogeny (98.4%), clustered apart from the *Agrobacterium* clade in the *recA* (73% similarity) and MLSA-based (76% similarity) phylogenies. In a previous study, the name *Rhizobium aggregatum* complex was proposed for the clade containing strain *A. albertimagni* AOL51^T^ [40]. However, in our study both the MLSA and WGS phylogeny and ANIb analysis showed that *A. albertimagni* AOL51^T^ is more closely related to *Allorhizobium*, suggesting that “*R. aggregatum* complex” is not the best choice. This is in agreement with Kuzmanovic et al. [55], who proposed the name “*Peteryoungia*” for strains formerly identified as “*R. aggregatum* complex”, and also included *A. albertimagni* AOL51^T^.

In the *Rhizobium* clade, *R. rhizogenes* TPD 7009 should be reclassified as *Agrobacterium* sp., *R. rhizogenes* Y79 and FIT62 should be reclassified as *Rhizobium* sp. and *Agrobacterium* strains 13-626, SHOUNA 12C, BETTINA 12B, and ICMP 7243 in the *Rhizobium rhizogenes* group should be reclassified as *R. rhizogenes*. Additionally, a set of strains identified as *Rhizobium*, *Allorhizobium*, or *Agrobacterium* clustered in three groups next to the *Agrobacterium* clade. These results are in agreement with [64], who suggested the transfer of these strains to the genera *Affinirhizobium*, *Alirhizobium*, and *Neorhziobium*. Similarly, strains identified as *Rhizobium* or *Agrobacterium* clustered also apart in three groups already suggested by Ma et al. [64] as *Peteryoungia*, *Paenirhizobium*, and *Heterorhizobium*.

### Rhizogenic strains are dispersed over at least two genera and at least nine (genomo)species

Another objective in this study was to shed more insight into the taxonomy of strains causing HRD. Interestingly, while HRD was originally associated with pathogenic *R. rhizogenes* (former biovar 2) strains, in recent studies it was observed that on (hydroponically grown) *Cucurbitaceae* and *Solanaceae* plants the causative agent is generally *Agrobacterium* biovar 1 harbouring a root-inducing plasmid [5, 6, 73, 107].

In previous research, Bosmans et al. [73] demonstrated that rhizogenic agrobacteria isolated from infested hydroponic greenhouses showed a remarkable phenotypic and genetic diversity. This genetic diversity is also seen here, since the rhizogenic strains included in this study are scattered over at least seven *Agrobacterium* (genomo)species (G2, G3, G7, G9, G19, G20, *A. cucumeris*, and *A. leguminum*) [73, 108]. Strain ST15.13.057, which was clearly separated from the other genomospecies, could be the first strain to be assigned to new genomospecies G23. This particular strain was isolated from tomato plants grown in a hydroponic greenhouse located in Belgium in 2014 [73], and showed high symptom severity compared to most other rhizogenic *Agrobacterium* strains [Kim et al. in prep]. This strain clustered closest to G8 strains, but only with 92.4-92.9% identity, supporting the idea that this strain could be a member of a new genomospecies. Nevertheless, more strains are needed to define the limits of this candidate new genomospecies and more biochemical analysis are required to confirm it as a new genomospecies [105]. Additionally, a G1 strain (ST15.13/045) and a G4 strain (KACC 21759) for which no genome sequences were available at the moment, were also reported previously to induce HRD [73, 108]. Although rhizogenic *Agrobacterium* strains are assigned to several genomospecies, most isolated *Agrobacterium* strains inducing HRD belong to G9 and to the bona fide species *A. cucumeris*. Nevertheless, so far no rhizogenic strains classified as G6 nor G13 were observed in our study, not ruling out that future isolations may detect rhizogenic strains in these genomospecies as well.

In contrast to rhizogenic *Agrobacterium* strains that were scattered over diverse genomospecies, the rhizogenic *Rhizobium* strains (previous *A. rhizogenes*) clustered closely together, including the type strain *R. rhizogenes* NBRC 13257^T^ (=LMG150^T^). Interestingly, the *R. rhizogenes* subclade also contained the avirulent *Rhizobium radiobacter* K84 strain (former *A. radiobacter* K84), which is known for its antagonistic activity towards *A. tumefaciens* causing crown gall disease in field crops [109]. The presence of *R. radiobacter* K84 within the *Rhizobium* clade in the three reconstructed phylogenies supports its reclassification into *R. rhizogenes* [22]. All these results highlight the fact that rhizogenic agrobacteria are polyphyletic, being located in at least two genera in the Rhizobiaceae family.

Finally, the rhizogenic strain *R. rhizogenes* YR147 and OV677, clustered as singletons closely to the *R. rhizogenes* subclade in all the phylogenies reconstructed and shared 90-91% identity according to ANIb with other *R. rhizogenes* strains, suggesting these strains needs a different specific epithet.

### Combination of MLSA and ANIb analysis is the best practice for robust classification into genomospecies in the *Agrobacterium tumefaciens* species complex

Compared to the *recA* tree, the MLSA and WGS phylogenies reflected better the genomospecies delineation provided by the ANIb groups, suggesting that the *recA* phylogeny, which has been initially used for genomospecies classification is not robust enough. The use of ANI has been suggested as a tool for resolving ambiguous phylogenies, which has become possible thanks to the enormous evolution in sequencing technologies [110]. ANIb has already been successfully used for accurate species delineation in complex genera, such as *Pseudomonas*, *Arcobacter*, *Stenotrophomonas,* and *Burkholderia* [111–114]. As a result, we applied ANIb in this study in order to shed more insight in the taxonomy of *Agrobacterium*, aiming to conciliate or resolve contradictory results in the different phylogenetic trees. For species delineation, the cut-off value of 94-96% for ANI has been recommended for prokaryotic organisms [115, 116]. More specifically, López-Guerrero et al. [29] recommended 95% similarity as a threshold for the delineation of species in the *Rhizobiaceae* family. As a result, a number of studies have applied the same cut-off value for the delineation of species in the *Rhizobiaceae* family [35, 102, 117, 118]. However, using a considerably large set of strains in this study (>400), we were able to fine-tune the cut-off to 94.0%. This allowed the identification of 24 distinct (genomo)species within the *Agrobacterium* genus. ANIb analysis also enabled delineation of three distinct species that clustered together in the MLSA- and WGS-based phylogenetic tree, namely *A. rubi*, *A. bohemicum*, and *A. rosae*, showing the usefulness of using this approach in a complementary manner with phylogenetic analysis based on MLSA or WGS. However, the ANIb analysis cannot be used as a stand-alone tool to infer phylogenetic relationships. For instance, the ANIb-based hierarchical tree, showed that, in contrast to MLSA and WGS phylogeny, that the *A. rubi-larrymoorei* clade clustered within the *Rhizobium* clade. This different clustering can be partially explained by the fact that pairwise distances values resulting from the ANIb analysis were clustered based on the specific algorithm known as “complete linkage” [119]. Moreover, ANI analysis does not apply an evolutionary model like phylogenetic analysis, and it is not considered an appropriate method for estimating genome relatedness among more divergent genomes, i.e., effective delineation of relationships at genus and family levels, for which AAI or phylogenomic treeing approaches are considered more appropriate [120].

In addition to calculating pairwise ANI distances, AAI distances were also calculated since it has been suggested that a 95% identity threshold could be applied for prokaryotic species delineation [121]. The AAI analysis did not provide a high-resolution species level delineation as compared to ANIb. Indeed, some of the separate ANIb groups were merged in the same AAI group when a cut-off of 95% was applied for AAI analysis, indicating that AAI was not able to delineate closely related (genomo)species within the Atsc. This is in agreement with the recommendation of using AAI comparisons for more distant relationships, e.g., for genus delineation [122]. Additionally, it was shown that GC content was strongly associated with the phylogenetic relationship present among the strains as observed in the MLSA tree. In other words, the more phylogenetically close *Agrobacterium* strains are, the more similar GC content values they display, again confirming the robustness of the MLSA analysis.

Altogether, we showed that using five genes in the MLSA, allowed a secure phylogenetic placement of AtSC strains. Nevertheless, in some cases the MLSA phylogeny could be improved by the use of ANIb.

The application of WGS-based phylogeny or phylogenomics is expected to yield more accurate phylogenies in contrast to other approaches using one or a limited set of genes [122]. This is not surprising since both methods use whole genomes, with more informative power and providing a better resolution for taxa delineation. Nevertheless, in our study, the subclades topology observed using WGS was largely similar to that observed in the MLSA phylogeny. The major downside of WGS phylogeny, however, seems to be the computing time that is needed. On desktop computer with 256 GB of storage memory, 16 GB of RAM memory, and 8 CPUs available, the WGS phylogeny analysis (i.e., the ML phylogeny reconstruction) for 200strains can take approximately 15 days, with the bootstrap value calculation taking a similar time, for a total of 30 days of continuous analysis. However, if computing power is moderate (e.g.: 4 GB RAM and 120 GB storage capacity), a combination of (*in silico*) MLSA and ANIb would be a fast (approx. four hours) and cost-effective method to unequivocally classify strains into genomospecies in the AtSC. This suggestion is in line with that of Kämpfer and Glaeser [123], who claim that MLSA represents an intermediate analysis between single marker and genome wide-based approaches to resolve phylogenetic resolution at the species level.

## CONCLUSION

WGS-based phylogeny presents a highly accurate phylogenetic reconstruction of AtSC strains. Nevertheless, conducting a WGS-based phylogeny reconstruction is a computationally demanding process. We propose conducting an MLSA in combination with ANIb analysis for accurate taxonomic resolution among AtSC strains needing a limited computational power.

## ACKNOWLEDGEMENTS

The resources and services used in this work were provided by the VSC (Flemish Supercomputer Center), funded by the Research Foundation – Flanders (FWO) and the Flemish Government.

## ABBREVIATIONS

AAI: Average amino acid identity
ANI: Average nucleotide identity
ANIb: ANI based on BLAST
ANIm: ANI based on MUMmer
Atsc: Agrobacterium tumefaciens species complex
AIC: Akaike’s information criterion
BIC: Bayesian information criterion
CDS: Coding sequence
CGH: Comparative genome hybridization
DDH: DNA-DNA hybridization
GBDP: Genome BLAST distance phylogeny
GC: Guanine-cytosine
GTRCAT: CAT General Time Reversible
HRD: Hairy root disease
ICSP: International Committee on Systematics of Prokaryotes
IJSEM: International Journal of Systematic and Evolutionary Microbiology
MLSA: Multi-locus sequence analysis
ML: Maximum Likelihood
NCPPB: National Collection of Plant Pathogenic Bacteria
NMDS: Nonmetric multidimensional scaling
LPSN: List of Prokaryotic names with Standing in Nomenclature
OGRI: Overall genome relatedness indices
pRi: Root-inducing plasmid
pTi: Tumor-inducing plasmid
TETRA: Tetranucleotide frequencies
WGS: Whole genome sequencing

## ACKNOWLEDGEMENTS

We would like to thank Professor Crauwels for sharing ideas, discussing, and giving advice on genome assembly methods, and Mss Alexa de Knijf for assistance with figures.

## FUNDING

This study was supported by the Agentschap voor Innoveren en Ondernemem (HBC.2017.0816) and KU Leuven internal funding (grant C14/19/074 and C3/19/047).

## Author’s contributions

PRV: Conceptualization, Investigation, Methodology, Data analysis, Writing – original draft. NK and MV: Data analysis, Writing - review & editing. SÁP: Investigation, Methodology, Data analysis, Writing – review & editing. HR: Supervision, Funding acquisition, Writing – original draft, review & editing, Validation. All authors read and approved the final manuscript.

## ETHICS DECLARATIONS

### Ethics approval and consent to participate

Not applicable.

### Consent for publication

Not applicable.

### Competing interests

The authors declare that they have no competing interests.

## DATA AVAILABILITY

The datasets generated and/or analysed during the current study are available in the NCBI nucleotide database and the accession numbers are listed in the supplementary Table S1.

## SUPPLEMENTARY MATERIAL

### Additional file 1

**Figure S1. Phylogenetic tree based on the 16S rRNA universal marker gene of *Agrobacterium*, *Allorhizobium*, and *Rhizobium***. A maximum likelihood phylogeny using as input a collection of 412*Agrobacterium*, 8 *Allorhizobium*, and 159 *Rhizobium* strains is shown. The phylogeny was built from the alignment the 16S rRNA gene (1,495 bp) using IQTREE. Bootstrap support value >70% based on 1,000 repetitions is represented by branches in bold. Leaves have been highlighted according to the defined 16S rRNA subclades. The colors in the outer rings indicate the 16S subclade or the genomospecies assigned to the *Agrobacterium* strains. Type strains are indicated in bold (see detailed annotation in online version https://itol.embl.de/shared/pme_bim). The scale bar represents the number of expected substitutions per site under the best-fitting model for the marker gene used (TN+F+I+G4 for 16S rRNA). Three *Ensifer adhaerens* strains (including type strain *E. adhaerens* Casida A^T^) were used as outgroup.

**Figure S2. Phylogenetic tree based on the *recA* marker gene of *Agrobacterium*, *Allorhizobium*, and *Rhizobium*.** A maximum likelihood phylogeny using as input a set of 412 *Agrobacterium*, 8 *Allorhizobium*, and 159 *Rhizobium* strains is shown. The phylogeny was built from the alignment the *recA* gene (1,089 bp) using IQTREE. Bootstrap support value >70% based on 1,000 repetitions is represented by branches in bold. Leaves have been highlighted according to genus affiliation. The outer rings show the ANIb subgroups assigned to the strais (STN: singleton), the defined MLSA and *recA* subclades as well as the genomospecies assigned to the *Agrobacterium* strains. Leaves highlighted in bold represent type strains (see detailed annotation in online version https://itol.embl.de/shared/pme_bim). The scale bar represents the number of expected substitutions per site under the best-fitting model for the marker gene used (TIM3+F+I+G4 for *recA*). Strain that did not cluster with any other are named as STN for singleton. Three *Ensifer adhaerens* strains (including type strain *E. adhaerens* Casida A^T^) were used as outgroup.

**Figure S3. Histogram of pairwise average nucleotide identity using BLAST (ANIb) values.** Pairwise ANIb values are shown for 412 *Agrobacterium*, 8 *Allorhizobium*, and 159 *Rhizobium* genomes. The pyani 0.2.10 package was used to calculate all pairwise ANIb values. The dashed red line represents this study’s optimized ANIb threshold for the delineation of *Agrobacterium* species at 94.0%.

**Figure S4. Principal component analysis (PCA) of normalized relative synonymous codon usage (NRSCU) obtained for different Atsc genomospecies.**(A) Scores plots showing the distribution of the Atsc genomes analyzed in this study according to the first two components (which explained 79.7% of the total variance) obtained by principal component analysis (PCA) of the normalized relative synonymous codon usage (NRSCU) values. Colors of points and the 68% data concentration ellipses denote different ANIb groups. (B) PCA loading plot, where each dot represents the loadings on the first two principal components for one factor (NRSCU for a particular codon).

### Additional file 2

**Table S1. List of genome assemblies downloaded from the NCBI database and sequenced in this study.**

**Table S2. List of markers used in this study and retrieved from the GCF_001649535 assembly for strain K599, also known as NCPPB2659.**

**Table S3. GC content of selected strains from the *Agrobacterium tumefaciens* species complex.**

**Table S4. Matrix presenting the pairwise values resulting from average nucleotide identity analysis using BLAST (ANIb).** The analysis was conducted for the 579 *Agrobacterium*, *Allorhizobium*, and *Rhizobium* quality-checked assemblies in this study. STN represents singletons and highlighted in orange, while 22 in-house sequenced rhizogenic *Agrobacterium* strains are highlighted in green.

**Table S5. Matrix presenting the pairwise values resulting from average amino acid identity (AAI) analysis.** The analysis was conducted among the 579 *Agrobacterium*, *Allorhizobium*, and *Rhizobium* quality-checked assemblies in this study.

## Notes

### Competing Interest Statement

The authors have declared no competing interest.

